# Characterization of epigenetic alterations in esophageal cancer by whole-genome bisulfite sequencing

**DOI:** 10.1101/2021.12.05.471340

**Authors:** Feng Pan, Shuai-Xia Yu, Xuan Wang, He-Cheng Huang, Zeng-Ying Cai, Jia-Min Wang, Shu-Yuan Lin, Yu-Lin Gao, En-Min Li, Li-Yan Xu

**Author notes:** Corresponding Authors: Dr. Li-Yan Xu, Shantou University Medical College, No. 22, Xinling Road, Shantou 515041, Guangdong, China.; and Dr. En-Min Li.

## Abstract

Esophageal carcinoma is a common and aggressive malignancy, and its patients have dismal clinical outcomes. The epigenetic dysregulation in both major subtypes, esophageal squamous cell carcinoma (ESCC) and adenocarcinoma (EAC), awaits further characterization. Here, we perform whole-genome bisulfite sequencing (WGBS) on a total of 43 esophageal cancer and normal samples, generating one of the largest WGBS datasets in this cancer to date. Focusing on hypomethylated regions in cancer, we show that they are associated with increased chromatin activity and enhancer RNA expression. Using this large collection of WGBS dataset, we reveal and validate novel clusters in both ESCC and EAC. We further identify specific molecular features in each cluster, with potential clinical implications. These data together advance our understanding of the epigenetic alterations in esophageal cancer and provide a rich resource for the research community of this disease.

## Introduction

Over 450,000 deaths annually worldwide are caused by esophageal carcinoma[1]. Esophageal squamous cell carcinoma (ESCC) and adenocarcinoma (EAC) are the two predominant histological subtypes with distinct geographical and epidemiological characteristics. ESCC predominates in the upper and mid-esophagus and is associated with smoking and alcohol consumption; EAC occurs near the gastric junction, and is associated with obesity, gastric reflux and Barrett’s esophagus. A third subtype, esophagogastric junction (GEJ) adenocarcinoma, strongly resembling EAC in terms of genomic, biological and clinical features, is typically treated similarly as EAC[2].

Although ESCC accounts for about 90% of esophageal cancers worldwide, the incidence of EAC has increased more than 8 fold in the western countries over the last 4 decades[3]. The prognosis for patients with esophageal cancer is poor in general: 5-year survival rate of EAC patients is approximately 17%, slightly higher than that of ESCC patients[4]. Oncological treatment options for esophageal cancer patients are limited and often ineffective, and surgical operations are associated with significantly deceased life quality. The molecular basis of both subtypes is still poorly understood, hindering the development of more effective therapeutic therapy. Despite numerous new insights gained from recent large-scale genomic analyses[2,5–14], these findings have not been translated into meaningful therapeutic strategies. Indeed, genome-driven targeted treatments for locally-advanced or advanced EAC individuals are few and largely ineffective. Moreover, targeted therapies have not even been established for ESCC except for immune checkpoint blockade therapy. Clearly, alternative molecular approaches at the non-genomic level are desperately needed to characterize the molecular pathogenesis of these two diseases for development of more innovative and effective regimens.

While these subtypes of esophageal cancers have major differences in genomic alterations, disparities at the levels of the transcriptome and DNA methylome[2,6] are also high. Indeed, DNA methylation profiling using Infinium 450k array identified widespread differentially-methylated regions between EAC and ESCC, and some of which were clearly associated with alterations in gene expression levels. For example, CDKN2A promoter region is much more frequently hypermethylated in EAC than ESCC[2]. However, much of DNA methylation work in esophageal cancer (and cancer in general) has focused on changes occurring at promoter regions, which are often hypermethylated in cancer cells[15,16]. This is because the majority of such investigations were based on methylation array profiling (e.g, Infinium HumanMethylation450 array), which by design covers much more densely gene promoter regions than the rest of the genome. While these array-based investigations have generated valuable knowledge in cancer epigenome, the single-nucleotide resolution of DNA methylation at the genome-wide level of esophageal cancer (and most other cancer types) remains poorly studied. A recent work compared 10 ESCC tumors with 9 nonmalignant esophageal squamous samples using whole-genome bisulfite sequencing (WGBS)[17]. However, the vast majority of DNA methylomes of EAC and ESCC have not been characterized, and in particular, the functional relevance of cancer hypomethylation in regulation of gene transcription is unclear.

To address these knowledge gaps, here we performed WGBS in a total of 43 samples from esophageal cancer, comprising both tumor and nonmalignant samples, and established methylome landscapes in both ESCC and EAC. We identified differentially methylated regions (DMRs) in cancer and revealed the epigenomic and transcriptomic impact of these methylation changes. Furthermore, we identified novel subtypes in both ESCC and EAC using WGSB data, and revealed distinct molecular features associated with these subtypes. These results together shed novel insights into the epigenetic regulation in esophageal cancer.

## Materials and Methods

### Whole genome bisulfite sequencing (WGBS) and data analysis

Primary tumors and nonmalignant samples were collected at Shantou center hospital, Medical College of Shantou University. The study of using human samples was approved by the ethics committee of the Medical College of Shantou University. Following DNA extraction and quality control (QC), 3 ug of DNA which was spiked with 26 ng lambda DNA were fragmented by sonication. Next, cytosine-methylated molecular barcodes were ligated to the fragmented DNA. Bisulfite conversion was performed using EZ DNA Methylation-GoldTM Kit (Zymo Research) according to the manufacturer’s instructions. DNA fragments were then PCR amplified by HiFi HotStart Uracil and ReadyMix (Kapa Biosystems). The clustering of the index-coded DNA samples was performed by Illumina cBot Cluster Generation System, followed by sequencing using Illumina Hiseq platform.

For WGBS data analysis, the sequencing reads were first aligned to the genome (build GRCh38) using the Biscuit pipeline (https://github.com/zwdzwd/biscuit). Duplicated reads were marked and removed by the Picard MarkDuplicates function (https://broadinstitute.github.io/picard/). We then measured the DNA methylation ratio using the Biscuit pipeline. CpGs with fewer than 3 reads of coverage were excluded from further analysis. QC was performed using TrimGalore for Illumina sequencing platforms (https://www.bioinformatics.babraham.ac.uk/projects/trim_galore/), PicardTools and MultiQC (https://multiqc.info/). Bisulfite conversion was measured by the Biscuit QC module in the MultiQC package (https://github.com/ewels/MultiQC/tree/master/multiqc/modules/biscuit).

To identify differentially methylated regions (DMRs), we first removed common PMD regions defined by Zhou et al. [18] (as shown in Fig.1B). Then the Dmrseq package was used to identify DMRs between tumor and nonmalignant samples using the following parameters: cutoff =0.1, bpSpan=1000, minInSpan=30, maxPerms=500[19]. Statistical cutoff of q value < 0.05 and absolute delta methylation change > 0.2 were applied for the DMR annotation.

**Figure 1.**
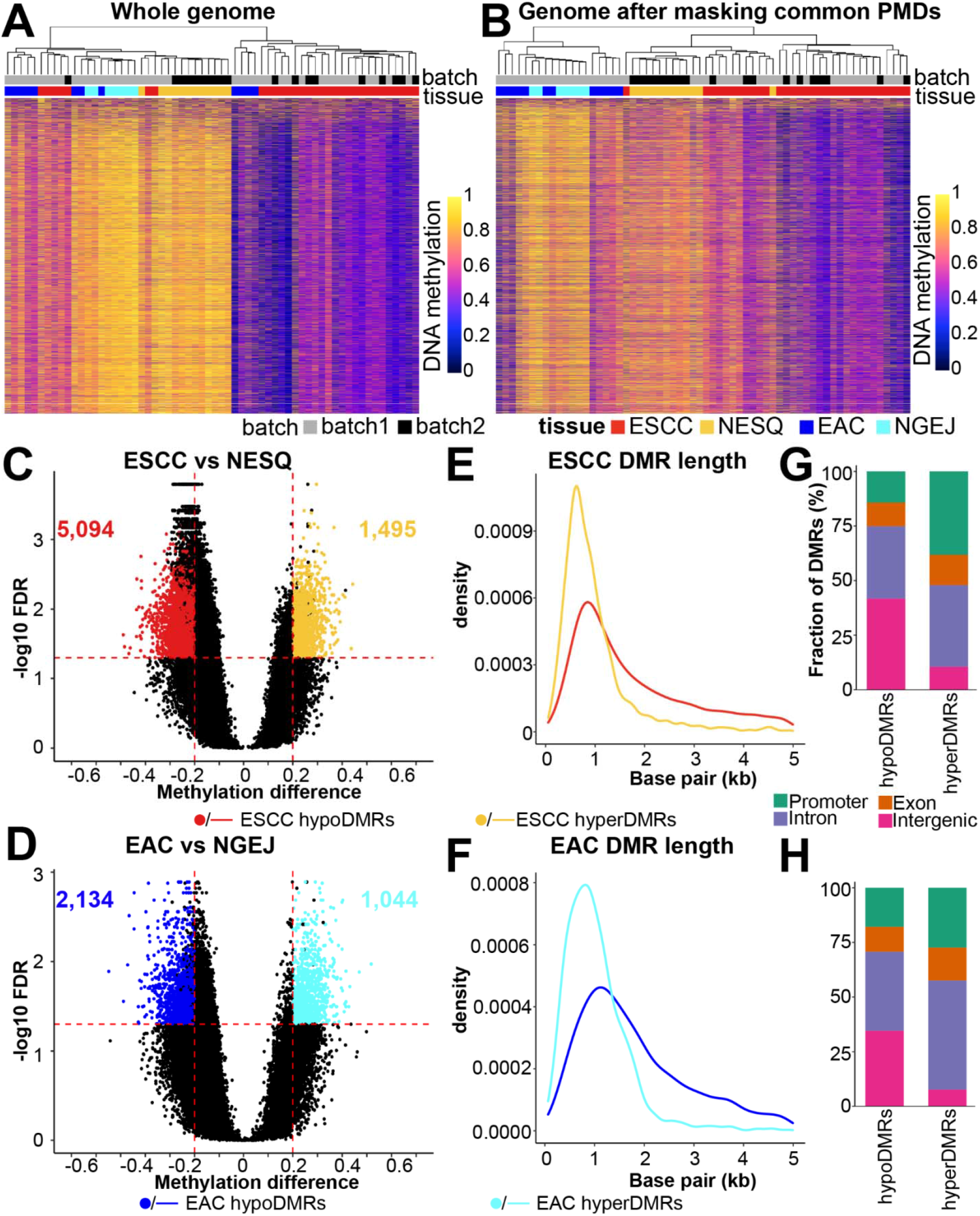
Methylome landscapes of esophageal tumors and nonmalignant samples. **(A-B)** Unsupervised clustering using top 8,000 most variable CpGs of all WGBS samples before **(A)** and **(B)** after removal of common PMD regions. Batch1 data was generated in this study, while Batch2 data was from Cao et al[17]. **(C-D)** DMRs were identified from the comparison between **(C)** ESCC vs. NESQ and **(D)** EAC vs. NGEJ, using cutoff as FDR < 0.05 and absolute delta methylation levels > 0.2. **(E-F)** Density plots showing the size of DMRs in **(D)** ESCC vs. NESQ and **(F)** EAC vs. NGEJ. **(G-H)** Stacked bar plots displaying fractions of DMRs in different genomic regions in **(G)** ESCC vs. NESQ and **(H)** EAC vs. NGEJ.

### Gene ontology (GO) term enrichment analysis by Cistrome-GO

The GO term enrichment analysis was performed using the Cistrome-GO method[20]. Specifically, hypoDMRs and upregulated genes in tumor compared with nonmalignant samples were used as the input data. The top 15 enriched pathways are displayed.

### Locus Overlap Analysis (LOLA)

The Locus Overlap Analysis (LOLA) was performed using the LOLA package[21]. Tumor hypoDMRs were used as the foreground sets and initial candidate regions identified by Dmrseq containing multiple methylation loci in all samples were used as the background sets. We used the default reference of genomic region sets as the database. The top 15 enriched terms with adjusted P value < 0.05 are displayed.

### Gene Set Enrichment Analysis (GSEA)

GSEA analysis was performed in a Preranked mode. For each molecular subtype, the fold change of gene expression was calculated compared to other remaining samples. We performed enrichment analyses in all expressed genes with mean FPKM values > 0.5 and used the expression fold change as the input and the Cancer hallmark gene sets as the database.

## Results

### Methylome landscapes of esophageal cancer and nonmalignant samples

To understand genome-wide DNA methylation alterations in esophageal cancer, we performed WGBS on 33 primary tumor samples, including ESCC (n=21), EAC (n=5) and GEJ tumors (n=7). We analyzed EAC and GEJ tumors together as one group given their strong similarity across genomic, biological and clinical characteristics (referred to as the EAC group for simplicity). Considering the distinct cell-of-origins of these two different subtypes, we used nonmalignant esophageal squamous epithelium (NESQ, n=3) and nonmalignant esophagogastric junction tissues (NGEJ, n=7) as controls for ESCC and EAC, respectively. We generated a mean of 10× sequencing depth per sample (**Supplementary Table 1**). Overall, > 75% of CpG dinucleotides with a coverage >=3 reads were detected in most samples. To increase the statistical power, we included the aforementioned WGBS dataset with 10 ESCC tumors and 9 NESQ samples[17], upon confirmation of the absence of batch effect (**Top annotation bar in Fig. 1A-B**).

To gain an overview of the methylomes across these samples, we performed unsupervised clustering using the top 8,000 most variable CpGs. However, this approach did not completely separate the two subtypes, with several ESCC and EAC samples mixed in the other subtype (**Fig. 1A**). We reasoned that global methylation loss in cancer might account for the mix-classification of some of the tumor samples. Indeed, one of the most pervasive methylation changes in cancer cells is the loss of DNA methylation globally in large hypomethylated blocks, which are termed partially methylated domains (PMDs). PMDs can cover more than one-third of the genome and are shared by different cancer types[22,23]. Therefore, the majority of the PMDs are likely shared between ESCC and EAC, and thus may mask specific methylation differences between ESCC and EAC. Considering this, we next removed common PMD regions using published annotations[18] and repeated the unsupervised clustering. Importantly, upon PMD removal, all squamous samples (ESCC and NESQ) were clustered together and completely separated from EAC and NGEJ samples (**Fig. 1B**). These results demonstrate that the most variably methylated CpGs outside PMDs represent cell-type-specific CpGs. Overall, these CpGs were more strongly methylated in nonmalignant samples than tumors, regardless of the cell type (**Fig.1B**).

We next determined differentially methylated regions (DMRs) between tumor and nonmalignant samples within each cancer type using Dmrseq package. A total of 5,049 and 1,495 DMRs were respectively hypomethylated (hypoDMRs) and hypermethylated (hyperDMRs) in ESCC (**Fig. 1C**), under the cutoff of q value < 0.05 and absolute delta methylation change > 0.2. Compared with ESCC, EAC had fewer hypomethylated DMRs (2,134) and a comparable number of hypermethylated DMR (1,044, **Fig. 1D**). Regardless of the cancer subtype, the majority of DMRs was about 0.5-2 kb long (**Fig. 1E-F**). Expectedly, in both cancer subtypes, tumor hyperDMRs were more enriched in the promoter regions and more depleted in intergenic regions than hypoDMRs (**Fig. 1G-H**).

### Epigenomic features of hypoDMRs in esophageal cancer

While hypermethylated regions in tumors have been extensively studied in many cancer types, including esophageal cancer[2,17,24], tumor hypoDMRs have received less attention since i) most of these changes cannot be captured in the 450K array, and ii) many of these regions fall into PMDs which are difficult to investigate. We therefore focused on studying this group of DNA methylation changes in cancer. We first determined H3K27ac levels in tumor hypoDMRs using ChIP-seq data from primary samples using either internal[25] or public datasets[26]. Notably, tumor hypoDMRs in both subtypes were associated with increased H3K27ac modification, indicating more active transcription regulation (**Fig. 2A-B**). We also analyzed enhancer RNA (eRNA) expression in tumor hypoDMRs using TCGA RNA-seq data[27]. Importantly, eRNA levels were much higher in tumor hypoDMRs in EAC than NGEJ (**Fig. 2C**). We were not able to analyze ESCC since only one nonmalignant squamous sample was available in TCGA data.

**Figure 2.**
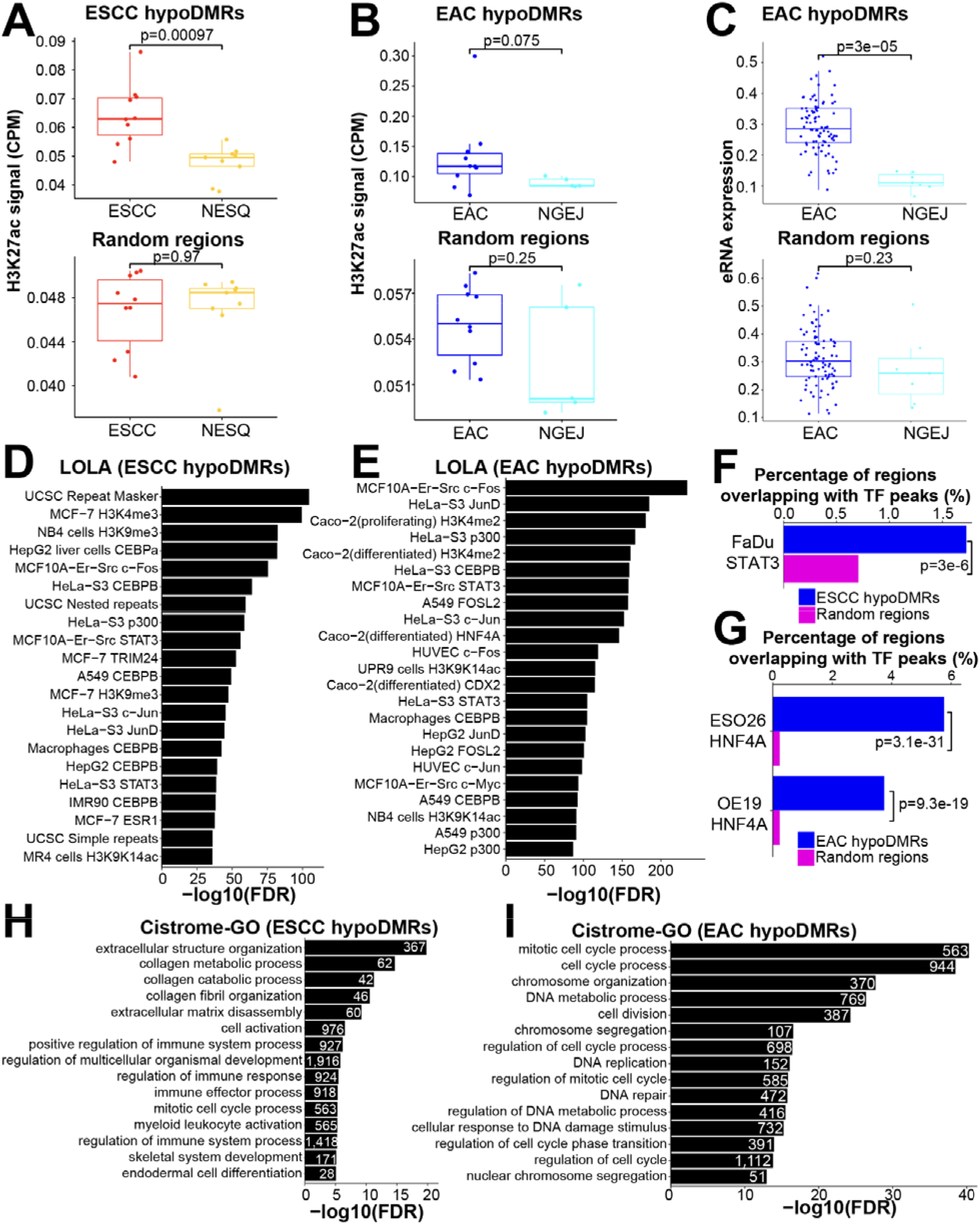
Epigenomic features of hypoDMRs in esophageal cancer. **(A-B)** Box plot showing H3k27ac signals in either **(A)** ESCC or **(B)** EAC hypoDMRs (upper) or random genomic regions (lower) using H3K27ac ChIP-seq data. **(C)** Box plots showing eRNA expression levels in EAC hypoDMRs (upper) or random genomic regions (lower) using TCGA RNA-seq data. **(D-E)** Enrichment of **(D)** ESCC or **(E)** EAC hypoDMRs in various epigenetic regions using the LOLA method. **(F-G)** Barplots showing fractions of regions overlapping with indicated ChIP-seq TF peaks in **(F)** ESCC or **(G)** EAC hypoDMRs. Note that Fadu is a cell line from head neck squamous cancer, which strongly resembles ESCC at biological and epigenetic levels. Fadu was used since public STAT3 ChIP-seq data in ESCC is not available. **(H-I)** GO term enrichment of genes associated with **(H)** ESCC or **(I)** EAC hypoDMRs using the Cistrome-GO method. The top 15 enriched pathways are shown; the numbers of enriched genes are indicated beside the bars.

We next investigated the distribution of DMRs along functional epigenomic domains, using the data from the Encode consortium[28]. As anticipated, in both cancer subtypes, tumor hypoDMRs were largely enriched in transcription factor (TF) binding sites and active chromatin domains, such as those marked with H3K4me3 and H3K4me2 (**Fig. 2D-E**). We did observe that repetitive sequences and H3K9me3 regions were also enriched in ESCC hypoDMRs, which could be due to incomplete removal of ESCC-specific PMDs. Most of the identified TFs were known to be active in their corresponding tumor types, such as STAT3 in ESCC[29,30], HNF4A[31,32] and CDX2[33] in EAC as well as AP1 factors in both cancer types. Using available ChIP-Seq data from our group (HNF4A ChIP-seq in ESO26 cells [32]) or others (HNF4A ChIP-seq in OE19 cells [31] and STAT3 ChIP-seq in FaDu cells [GSE78212]), we further validated that binding peaks of these TFs were significantly enriched in tumor hypoDMRs (**Fig. 2F-G**).

We next explored potential biological significance of tumor hypoDMRs in esophageal cancer by using genes associated with hypoDMRs based on the Cistrome-GO method[20]. Known pathways and cellular processes important for ESCC were identified, including “extracellular structure organization”, “extracellular matrix disassembly”, “mitotic cell cycle process”, etc (**Fig. 2H**). In EAC, we found that most pathways were related to cell cycle progression and DNA replication (**Fig. 2I**).

### DNA methylome analyses identify molecular subtypes of ESCC

Subclassification of cancer patients based on molecular features greatly improves traditional histopathological taxonomy and helps establish precision medicine. ESCC is known to have strong inter-tumoral heterogeneity, however, the classification and subtyping of ESCC patients await further improvement[2,34,35]. Previously, DNA methylation profiling has been utilized to successfully establish important cancer subtypes, such as those in colon cancer[36], gastric cancer[37], medulloblastoma[38], etc. Therefore, we next sought to identify ESCC subtypes using WGBS data.

Performing an unsupervised clustering approach using the top 8,000 most variable CpGs, we identified three clusters in ESCC, independent of the sequencing batch (**Fig. 3A**). At the DNA methylation level, Cluster-1 showed an overall hypermethylated pattern (**Fig. 3B**). We next investigated TCGA ESCC samples as an independent cohort, using overlapping probes on the 450K methylation array with the most variable WGBS CpGs. Importantly, we validated the presence of the three clusters in TCGA samples **(Fig. 3C)**, with highly consistent ratios between these three clusters (**Fig. 3D**). We further interrogated genomic alterations between these clusters using the TCGA data, focusing on significantly mutated genes (SMGs) as annotated by prior research [2,13,14,39]. The two most frequently altered drivers, TP53 and CDKN2A, showed comparable mutation/deletion rates across the three clusters (**Fig. 3E**). Interestingly, Cluster-1 exhibited higher alteration rates in tumor suppressor genes including ZNF750 and FAT1, while Cluster-2 had increased genomic changes in TP63, NFE2L2, PTCH1 and CUL3. These data demonstrate that the three ESCC clusters are driven by different genomic lesions, which influence distinct pathways. To further corroborate this observation, we determined differentially expressed genes between the three clusters (**Fig. 3F**) and performed Gene Set Enrichment Analysis (GSEA) enrichment analysis using the Cancer Hallmark gene set. Supporting that Cluster-2 had higher NFE2L2 and CUL3 mutations, which regulate oxidation and antioxidant processes, “Reactive oxygen species pathway” and “Oxidative phosphorylation” were strongly and exclusively enriched in this cluster **(Fig. 3G)**. In Cluster-1, all top 5 enriched pathways were immune related, including “Allograft rejection”, “Interferon alpha response”, “Inflammatory response”, “Interferon gamma response” and “IL6_JAK_STAT3 pathway”, suggesting that this subtype has stronger active immune response and possibly higher immune cell infiltration. In Cluster-3, only one pathway, “Angiogenesis” was significantly enriched.

**Figure 3.**
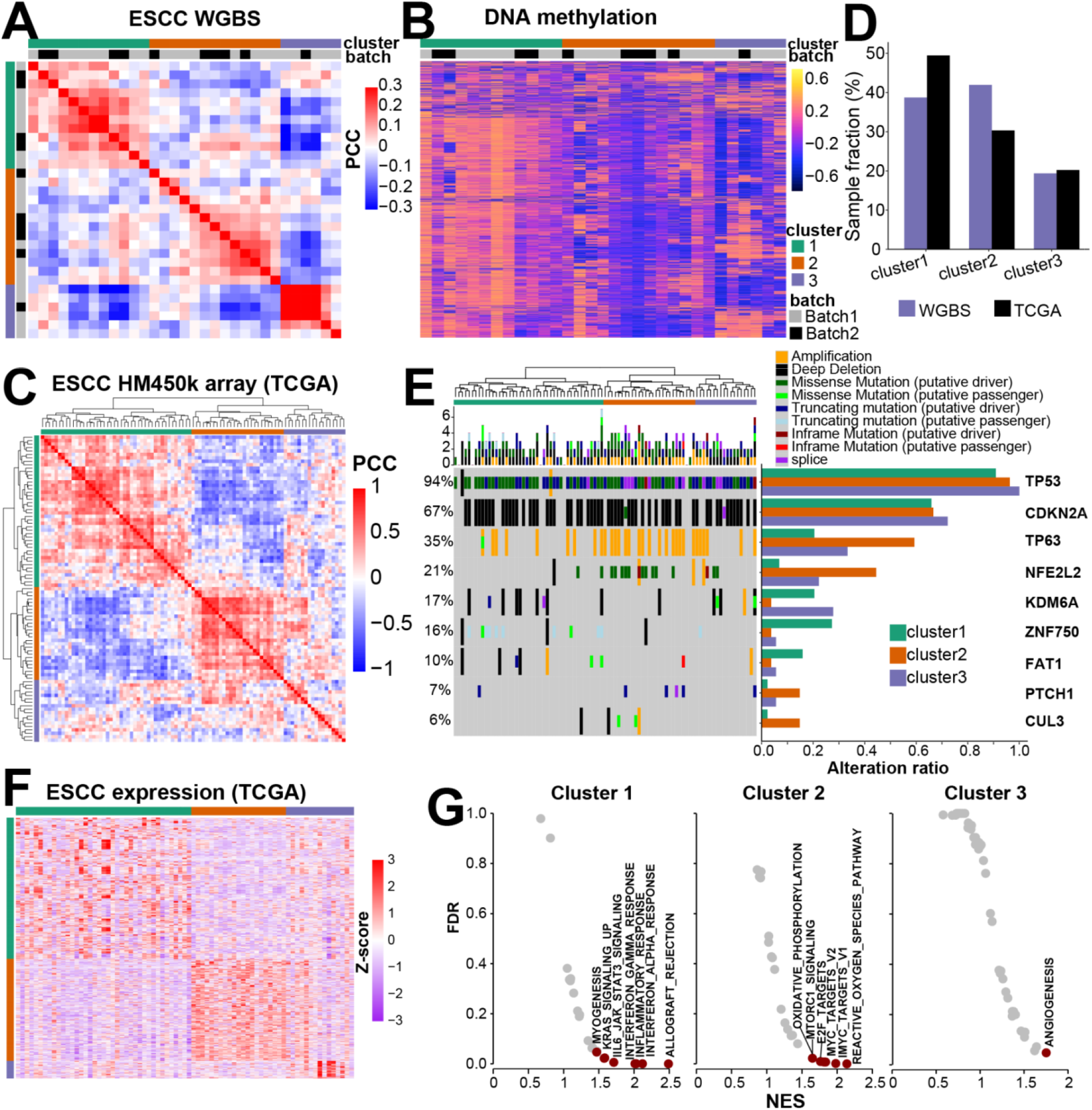
Molecular clustering of ESCC tumor samples using WGBS data. **(A)** A heatmap showing the Pearson correlation coefficient (PCC) among ESCC tumors profiled by WGBS. Cluster and batch information are annotated. **(B)** A heatmap displaying differentially methylated CpGs between the three clusters identified in **(A). (C)** A heatmap showing the Pearson correlation coefficient among TCGA ESCC tumors profiled by 450K methylation array. **(D)** The fraction of samples in each cluster in the two datasets. **(E)** An oncoplot showing genomic alterations in indicated driver genes, with barplots (right) displaying aggregated frequencies in different clusters. **(F)** A heatmap showing differentially expressed genes between the three clusters using TCGA RNA-seq data. **(G)** Dotplots showing enriched pathways in the three clusters using GSEA analysis based on the Cancer Hallmark gene sets.

### Two molecular subtypes of EAC

We similarly performed an unsupervised clustering analysis in EAC tumors and identified two clusters (**Fig. 4A**). These 2 clusters contained both EAC and GEJ tumors, as expectedly. At the DNA methylation level, Cluster-2 was overall more highly methylated compared with Cluster-1 **(Fig. 4B)**. Again, we validated that TCGA EAC samples were also separated into 2 clusters **(Fig. 4C)**, with a highly consistent ratio **(Fig. 4D)**. At the genomic level, while TP53 mutations were evenly distributed in these two clusters, Cluster-2 displayed higher alteration frequencies in receptor tyrosine kinase (RTK) pathway genes (ERBB2, VEGFA, KRAS) and Cluster-1 had more mutation/deletion rates in tumor suppressors SMARCA4 and APC **(Fig. 4E)**. In addition, ARID1B was exclusively mutated in Cluster-2. GSEA analysis using differentially expressed genes between these two clusters (**Fig. 4F**) revealed that cell cycle pathways (“G2M checkpoint” and “E2F targets”) were enriched in Cluster-2 (**Fig. 4G**), likely owing to higher activation of RTK signaling as indicated from the mutational analysis. On the other hand, immune related pathways were enriched in Cluster-1, including “Interferon gamma response”, “Allograft rejection”, “Interferon alpha response”, “Inflammatory response”, “IL6_JAK_STAT3 pathway” and “Complement”. Notaly, Cluster-2 patients had significantly better prognosis compared with Cluster-1 **(Fig. 4H)**, further suggesting that these two clusters have different biological and clinical features.

**Figure 4.**
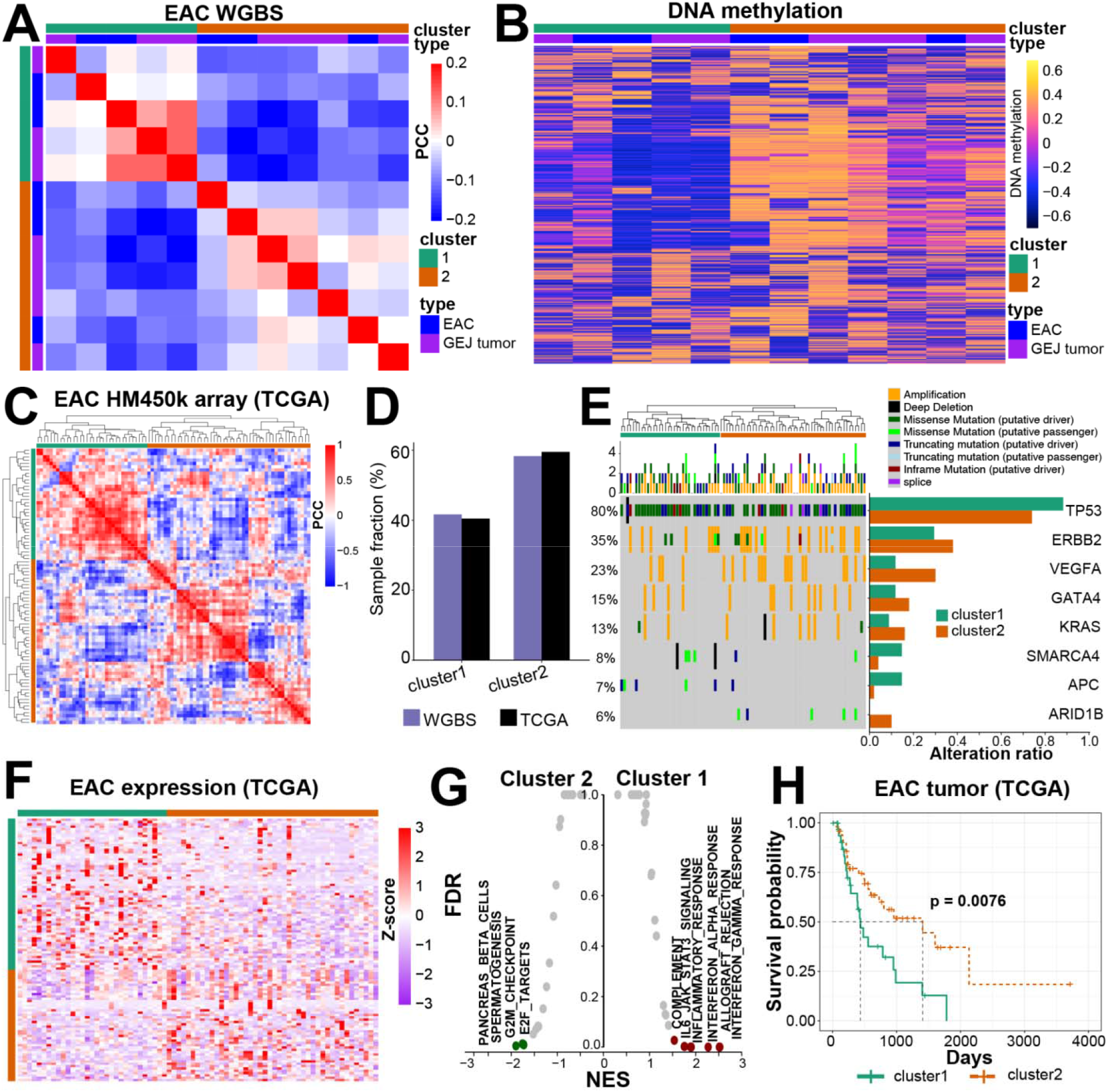
Molecular clustering of EAC tumor samples using WGBS data. **(A)** A heatmap showing the Pearson correlation coefficient among EAC tumors profiled by WGBS. Cluster and batch information are annotated. **(B)** A heatmap showing the differentially methylated CpGs between the two clusters identified in (A). **(C)** A heatmap showing the Pearson correlation coefficient among TCGA EAC tumors profiled by 450K methylation array. **(D)** The fraction of samples in each cluster in the two datasets. **(E)** An oncoplot showing genomic alterations in indicated driver genes, with barplots (right) displaying aggregated frequencies in different clusters. **(F)** A heatmap showing differentially expressed genes between the two clusters using TCGA RNA-seq data. **(G)** Dotplots showing the enriched pathways in the two clusters using GSEA analysis based on the Cancer Hallmark gene sets. **(H)** A Kaplan-Meier plot showing the survival probability of the two clusters of TCGA EAC samples.

## Discussion

In the present study, we generated and analyzed genome-wide methylation profiles of one of the largest cohorts of esophageal cancer at the single base resolution to date, from both ESCC and EAC patients along with the matched nonmalignant tissues. We augmented the DNA methylation results with genomic, transcriptomic, and chromatin modification data from external studies, providing integrative analyses of the epigenetic regulation in this malignancy. ESCC and EAC have distinct cell identity and gene regulatory programs, and we analyzed these two subtypes separately, using their corresponding nonmalignant samples. Indeed, the most variably methylated CpGs reflect the differences between cell types rather than changes between cancer and noncancerous tissues (**Fig. 1B**). These top variable CpGs are likely located within cell-type-specific regulatory elements (e.g., enhancer, promoters, insulators) that distinguish squamous and columnar/glandular cells. Target genes of these elements may contribute to the differentiation and cell-type-specific functions of these cell types.

In both ESCC and EAC, thousands of regions were differentially methylated between tumor and nonmalignant samples, indicating widespread DNA methylation rewiring during cancer development and progression. Regions that gain methylation (i.e., hyperDMRs) have been extensively investigated. Indeed, hypermethylated CpG island promoters in cancer, the most prominent cancer-specific epigenetic dysregulation identified decades ago, fall into this category. In contrast, regions that lose DNA methylation (i.e., hypoDMRs) in cancer have been given much less attention owing to the lack of representative probes on the 450K methylation array, the most commonly used platform for DNA methylation studies. Another important reason for the understudy of cancer hypoDMRs is that many of these regions are within PMDs which are enriched in heterochromatin and repetitive regions, and their direct biological significance in cancer can be obscure and difficult to understand. To overcome this challenge, we masked common PMD regions before DMR analysis. This approach successfully identified focal regions with methylation loss which were associated with active chromatin, increased eRNA expression and cancer-promoting pathways (**Fig. 1-2**). Nevertheless, methylation changes in large domains such as PMDs can also have functional impact in cancer biology, which warrants future investigations[18,40].

Molecular subclassification of cancer has imperative basic and translational values. For example, extensive investigations of breast cancer subtypes (e.g., luminal, basal-like, Her2) have revealed novel biological insights and led to transformative clinical management of its patients. Similarly, precision medicine has been applied to different subtypes in lung cancer such as EGFR-mutant and ALK-fusion tumors. However, molecular taxonomy of either ESCC or EAC patients remains comparatively less well-studied, despite the recognition of the presence of strong inter-tumoral heterogeneity in both cancer types. Based on our WGBS data, we identified three clusters in ESCC and two in EAC patients. Importantly, the existence of these clusters were validated in the TCGA cohort, which showed highly consistent ratios between these clusters. In ESCC, Cluster-1 was characterized by DNA hypermethylation, high mutation burden of KDM6A, ZNF750 and FAT1, as well as enriched immune pathways. Patients from this cluster may exhibit differential sensitivity to immunotherapy. ESCC Cluster-2 had more mutations targeting NFE2L2 and CUL3, both of which are key regulators of oxidative phosphorylation and redox reactions. Indeed, these pathways were strongly enriched in Cluster-2, highlighting metabolic stress in this subgroup of samples. In EAC, the two clusters not only differed in their driver mutation patterns and gene expression programs, but also showed distinct survival outcomes. Although further investigations are required to confirm and understand these clusters, these data highlight the existence of extensive inter-tumoral heterogeneity among esophageal cancers with distinct biological and clinical characteristics.

## Acknowledgements

This work was supported by grants from the National Cohort of Esophageal Cancer of China (2016YFC0901400) and 2020 Li Ka Shing Foundation Cross-Disciplinary Research Grant (2020LKSFG07B).

